# A framework for analyzing *C. elegans* neural activity using multi-dimensional hyperbolic embedding

**DOI:** 10.1101/2021.04.09.439242

**Authors:** Iulia Rusu, Zachary T. Cecere, Javier J. How, Kathleen T. Quach, Eviatar Yemini, Tatyana O. Sharpee, Sreekanth H. Chalasani

## Abstract

Neurons represent changes in external and internal environments by altering their activity patterns. While coherent brain-wide patterns of neural activity have been observed in neuronal populations, very little is known about their dimensionality, geometry, and how they are correlated with sensory inputs. Here, we recorded the activity of most head neurons in *Caenorhabditis elegans* experiencing changes in bacterial or control buffer stimuli around their nose. We first classified active neurons into six functional clusters: two sensory neuron clusters (ON and OFF responding to addition and removal of stimuli, respectively) and four motor/command neuron clusters (AVA, RME, SMDD and SMDV). Next, we estimated stimulus selectivity for each cluster and found that while sensory neurons exhibit their maximal responses within 15 seconds, changes in bacterial stimuli drive maximal responses in command and motor neuron clusters after tens of seconds. Furthermore, we show that bacterial stimuli induce neural dynamics that are best described by a hyperbolic, not Euclidean, space, of dimensionality eight. The hyperbolic space provided a better description of neural activity than the standard Euclidean space. This space can be separated into three components – one sensory, and two motor directions (forward-backward and dorsal-ventral). Collectively, we show that *C. elegans* neural activity can be effectively represented in low-dimensional hyperbolic space to describe a sensorimotor transformation.

**Significance statement:** A major function of a nervous system is to transform sensory information into behavioral outputs. As the first receiver of sensory input, sensory neuronal activity is often most correlated with stimulus features. However, this sensory activity is modified as it travels to other neurons, where it integrates with network activity before altering motor neurons and driving corresponding behavior. Activity in non-sensory neurons is driven by ongoing network activity and sensory input, but distinguishing between their relative contributions is often difficult. Here, we identify two sensory and four command/motor neuron clusters in the *C. elegans* neural network responding to bacterial stimuli and define their receptive fields. We then use a hyperbolic embedding to identify how these clusters interact with each other and identify the relevant dimensions that might alter behavior. Our method is fully scalable to other systems, including those without neuronal identities, and allows us to attribute neural activity to network states and behavioral outputs.

## Introduction

Animals are constantly adapting to changes in their environment. They use their sensory neurons to extract relevant information, process that information using various neuronal populations, and then drive motor behavior by altering motor neuron activity. One approach to understanding how sensory information is transformed into behavioral outputs has been to trace the changes in activity from sensory to motor neurons. Sensory neuronal responses are most correlated with stimulus features [1,2], but mechanisms including attention [3,4], cognitive load [5], learning [6,7], internal noise [8,9] and internally coordinated activity [10] can introduce variability. Additionally, sensory neurons are also gated by behavior, such as locomotion [11,12] and whisking [13]. On the other hand, some neuronal populations, including interneurons and motor neurons, are often poorly correlated with sensory information. Progress in understanding the brain has been limited as less is known about how neuronal populations interact with each other and how these interactions are altered by sensory activity.

The transparent nematode *Caenorhabditis elegans* with its small, well-defined nervous system is ideally suited to reveal insights into how an intact nervous system processes sensory information. These efforts have been aided by advances in microscopy [14], genetically encoded sensors [15], and transgenic animals where neurons can be unambiguously identified by the expression of different fluorophores (NeuroPAL, [16]). Additionally, studies have shown that calcium dynamics in neuronal nuclei mirror those observed in their cytoplasm [17], which means that a large fraction of nervous system activity (189 out of the total 302 neurons [18]) can be measured from a small region in the head of the animal. Previous efforts to analyze these whole-brain data used principal component analysis (PCA) to demonstrate that activity in this network switches between neurons driving forward, backward, ventral, or dorsal turns [19], and that this activity was altered by animals that were either food-deprived [20] or in a sleep-like state [21]. Other studies monitored neural activity in freely moving animals and found that these data could be analyzed using PCA to obtain interpretable results, showing forward and backwards locomotion across multiple sets of neurons [22]. More recent efforts have focused on biogenic amine circuits [23], or used optogenetic activation to monitor spread of neural activity across the network [24]. Despite these advances, it is still unclear how neuronal populations within this network work together to process sensory information and select the appropriate behaviors.

Here, we monitor the calcium activity of neurons in the head of the animal while its nose experiences a rapidly fluctuating sequence of bacterial stimuli. These stimuli are ethologically relevant to the animal as it feeds on the same strain of bacteria. We find that active neurons can be classified into six clusters (two each of sensory, inter-, and motor neuron classes). Next, we identified the names of neurons in the interneuron and motor neuron clusters by leveraging NeuroPAL transgenic animals. We then used maximum noise entropy (MNE) [33, 34] to define stimulus selectivity for each of these six clusters and found that each cluster responds to different features of the stimulus. Finally, we use a hyperbolic structure to show that this network is hierarchical and define how sensory stimulus alters the underlying dimensionality.

## Results

### *C. elegans* neurons can be classified based on their responses to bacterial stimuli

To monitor responses of *C. elegans* head neurons, we trapped transgenic animals, expressing the pan neuronal nuclear-localized genetically encoded calcium indicator GCaMP5K, in a transparent PDMS-based microfluidic device [17, 25]. We previously used this setup to reveal changes in neural network topology while the animals experienced various chemosensory repellents and attractants [26]. In this study, we instead delivered pulses of bacterial stimuli to the nose of the animals (Fig. 1a); we refer to this as the bacteria-buffer stimulus sequence. To control for artifacts intrinsic to the microfluidic setup, we also imaged activity while *C. elegans* was presented with a control buffer-to-buffer stimulus sequence that alternated between two flows composed of chemically identical buffer (**Fig. 1a**). We also recorded responses from animals expressing GCaMP6s along with several other transgenes that allowed for neurons to be unambiguously identified using a combination of fluorophores (NeuroPAL, [27], **Fig. 1b-d**). Data obtained from this strain was used to confirm cell identity associated with activity patterns found in the primary strain (**Fig. S1a**). The normalized calcium traces using min-max scaling, from a single animal experiencing a bacteria-to-buffer stimulus pattern are shown in **Fig. 1f**. We identified sensory neurons as those that were either positively (ON cells) or negatively (OFF cells) correlated with each of the bacterial pulses. Specifically, ON cells were those neurons whose activity increased upon all onsets of bacterial stimuli and decreased immediately upon removal of bacterial stimuli (Fig. 1g, green trace). Similarly, OFF cells had the opposite response – their activity increased upon removal of bacteria and decreased upon addition of bacteria (**Fig. 1g, orange trace**).

**Figure 1.**
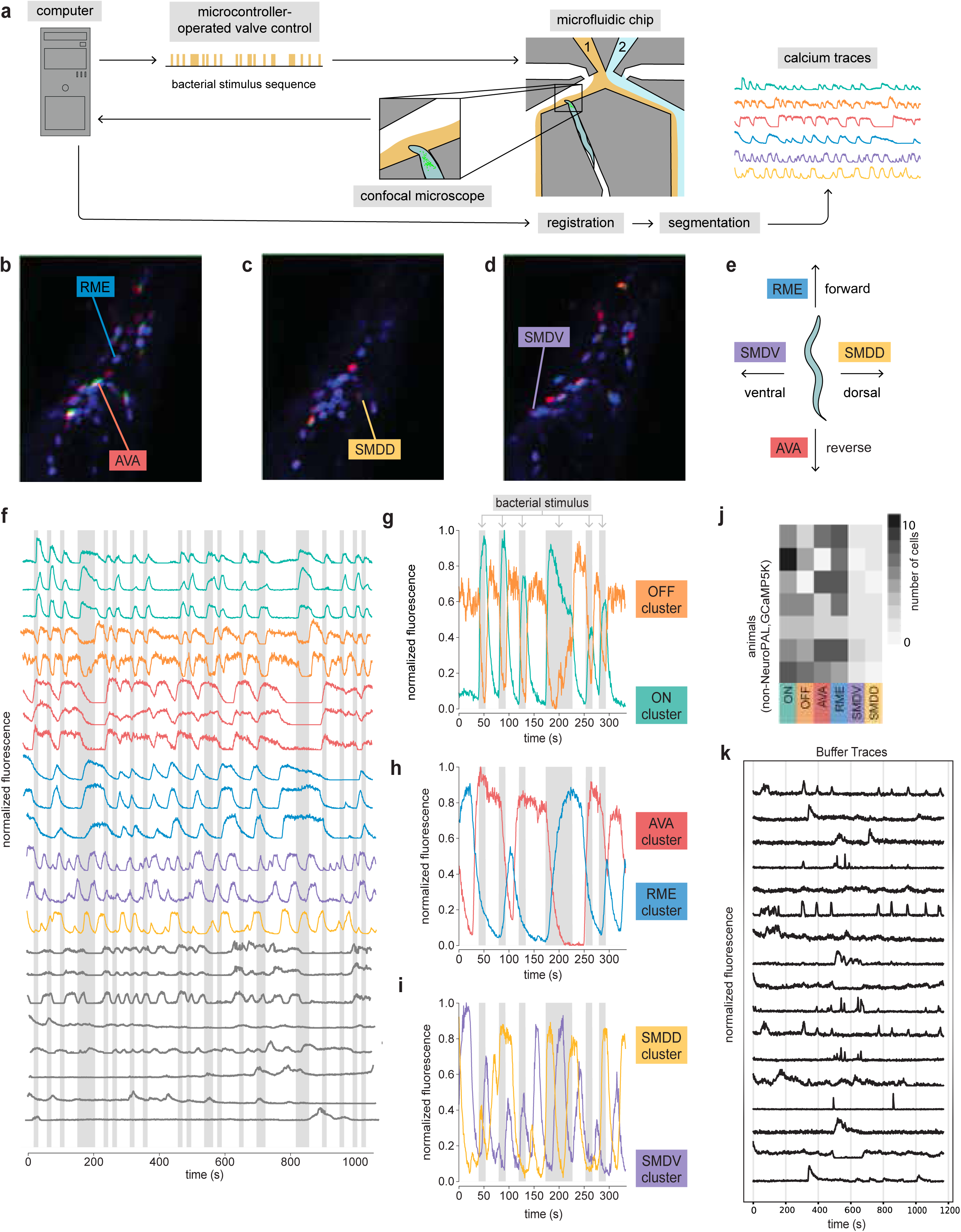
*C. elegans* neurons can be classified based on their responses to bacterial stimuli. (**a**) Stimulus presentation and imaging setup. A computer precisely controls the delivery of a stimulus sequence that alternates between variable-length pulses of two liquid flows: bacterial food stimulus (gold) and control buffer (light blue). This stimulus sequence is presented to the nose of a *C. elegans* that is restrained in a microfluidic chip. Volumes of the *C. elegans* head are acquired and subsequently processed to acquire changes in fluorescence, a proxy for intracellular calcium. (B–D) Identification of neurons in the NeuroPAL–GCaMP6s strain. Examples of (**b**) RME and AVA neurons, (**c**) SMDD and (**d**) SMDV neurons identified using NeuroPAL markers. (**e**) Neurons and their associated direction of locomotion. (**f**) Calcium activity of low-noise active neurons for a single worm. Gray shading represents bacterial stimulus pulse duration. Neurons are color-coded to one of six clusters (ON, OFF, AVA, RME, SMDD, SMDV), or grey if not assigned a cluster. (**g**) OFF and ON clusters. (**h**) AVA and RME clusters. (**i**) SMDD and SMDV clusters. (**j**) Number of neurons in each functional cluster for all animals, based on > 85% Pearson correlation with representative neurons. (**k**) Representative calcium traces from neurons in animals exposed to buffer alone.

Apart from sensory neurons, we were interested in identifying other subsets as well. We found a second pair of neurons whose activity was anti-correlated with each other and had prolonged time courses that were weakly correlated with stimulus onset (**Fig. 1h**). Using the NeuroPAL–GCaMP6s strain, we identified these neurons as the RME motor neurons and AVA command neurons. Anti-correlated and relatively long-lasting responses in these neurons have been previously observed in animals experiencing changes in ambient oxygen levels [19]. We also identified two other pairs of anti-correlated motor neurons – SMDD and SMDV – that exhibited moderately faster dynamics compared to the AVA/RME pair (**Fig. 1i**). These four pairs of neurons are associated with forward (RME) and backward (AVA) movements, and dorsal (SMDD) and ventral (SMDV) turns [19,28,29,30,31]. Motivated by the knowledge of multiple AVA and RME neurons existing in a single organism, we next sought to assign the remaining non-sensory neurons to the RME and AVA clusters based on a strong Pearson correlation (>85%) with the identified RME and AVA neurons, respectively (**Fig. 1h**). In contrast, SMDV and SMDD were often the sole members of clusters within each organism (**Fig. 1j**). Furthermore, we also found additional neurons whose activity pattern did not match any of these six clusters (**Fig. 1f**, grey). Notably, these unclustered neurons did not respond when we transitioned from buffer to buffer (**Fig. 1k**) We found activity patterns matching all clustered neurons, including ON and OFF sensory (**Fig. S1b**), SMDV and SMDD (**Fig. S1c**, R ∼ 0.4), AVA and RME neurons (**Fig. S1d**, R ∼ 0.5) in animals experiencing buffer-buffer sequences. Collectively, these data suggest that *C. elegans* neurons can be classified into at least six clusters (sensory ON and OFF neurons, AVA, RME, SMDD, and SMDV).

### Patterns of bacterial stimuli drive responses in the six neuronal clusters

To characterize stimulus transformation performed by neuronal clusters, we used a model that makes minimal assumptions about associations between stimuli and neural responses known as the maximum noise entropy model [32,33]. Here, neural responses are modeled by a sigmoidal nonlinearity (which is a property of least constrained neural response model):

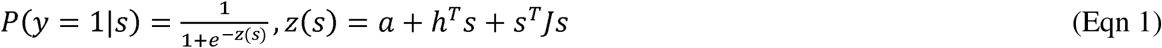

Here 𝑦 is the calcium response, 𝑠 is the stimulus, and we fit for 𝑎, ℎ, and 𝐽, which collectively determine the degree of the nonlinear response. We trained our model and extracted a low rank matrix 𝐽 that was determined by minimizing the negative log-likelihood [34]. The filter ℎ and matrix 𝐽 together describe the preferred stimulus features for each neuronal cluster.

The dominant preferred stimulus features of the six neuronal clusters are shown in **Fig. 2**. In these figures the x-axis indicates amount of time in seconds, till response onset, with zero indicating the response. We highlight each neuron’s temporally sensitive region in grey. In the inset of each figure panel, we plot the neural response as a function of the strength of the dominant stimulus features, given by projecting stimuli onto it. We found that ON and OFF sensory neurons typically had their maximal responses within 15 seconds of experiencing an increase or decrease in bacterial stimuli, respectively (**Fig. 2a, b**). This time window is consistent with previous reports that maximal neural activity in AWC, ASI, ASK and other chemosensory neurons were observed within 10 to 20s of the addition or removal of bacterial stimuli [35,36,37].

**Figure 2.**
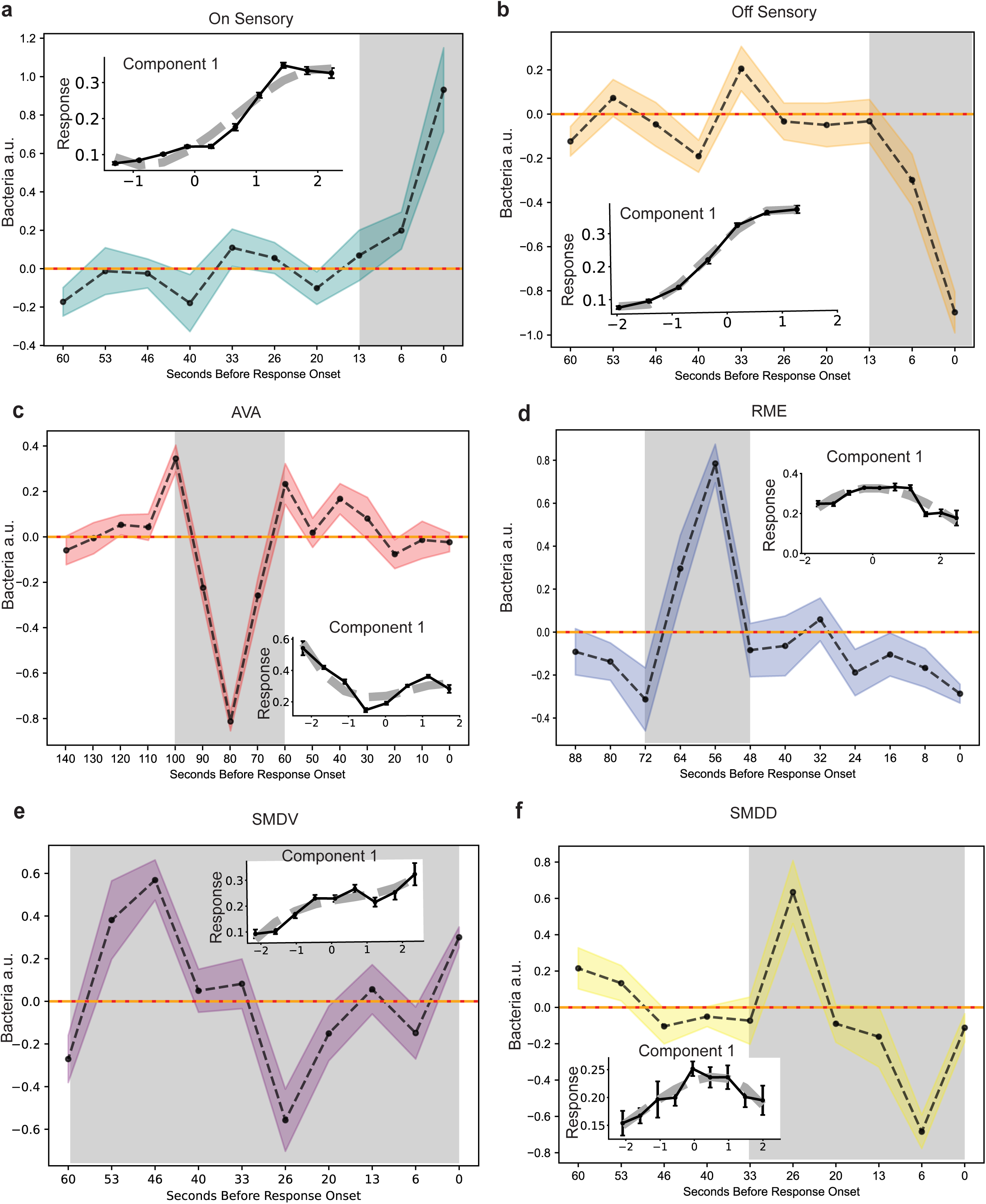
Patterns of bacterial stimuli drive responses in the six neuronal clusters. (**a**-**f**) Dominant preferred stimulus features of ON/OFF sensory, AVA, RME, SMDV, and SMDD clusters. Y-axis indicates presence or absence of bacteria, x-axis indicates time in seconds, before neural response. In each central panel, the black dashed line indicates the average eigenvector across 4 jackknives. The upper bound and lower bounds of the colored region are standard error of the mean, mean ± SEM, taken from 20 bootstraps. Highlighted in grey, is each neuron’s temporally sensitive region. Each inset is a plot of normalized calcium neural response as a function of the strength of the dominant stimulus features. Black solid line plotted to have zero mean and unit variance, error bars, mean ± SEM. Grey dashed lines represent a line of polynomial fit. All models were trained across data from 7 animals.

By comparison, interneuron AVA and RME clusters exhibited more complex types of stimulus selectivity. First, these interneurons were most sensitive to stimulus fluctuations that occurred several tens of seconds prior to their responses. Second, the responses of these neurons were quadratic with respect to the strength of these fluctuations (see insets in Fig. **2c** and Fig. **2d**) [36]. In other words, in each case neurons were similarly affected by the stimulus fluctuations regardless of their direction. However, whereas AVA neurons were activated by the fluctuation (**Fig. 2c, inset)**, highest response is seen at negative and positive stimulus in the x-axis. The RME neurons were suppressed by a fluctuation with a similar time course. (**Fig. 2d, inset)**, highest response is seen at values corresponding to zero stimulus on the x-axis. To understand the stimulus selectivity of both AVA and RME interneurons it was necessary to consider additional stimulus features (**Fig. S2**) that break the symmetry between different signs of stimulus fluctuations. For example, in the case of the AVA neuron, the secondary stimulus feature indicates that its responses were increased by bacterial removal (**Fig. S2g**) in addition to being strongly triggered by a bacterial fluctuation described by the primary stimulus feature. This type of selectivity is consistent with previous studies demonstrating that removing animals from bacteria has been shown to increase reversals, and which are in turn driven by AVA activity[38,39]. In the case of the RME interneuron, its secondary feature corresponded to an increase in bacterial concentration (**Fig. S2h**). Thus, the combined primary and secondary feature selectivity for this interneuron indicate that it is activated by fluctuations that indicate the introduction of bacteria. This is also consistent with other previous studies where RME interneuron was shown to drive forward motion.

The activity of motor SMDD and SMDV neurons was simpler than that of interneurons and can be explained largely by the primary stimulus feature alone. Specifically, SMDD neurons were most active when the animal experienced oscillatory patterns with medium length scales, over 30 seconds. By comparison, SMDV neurons were sensitive to longer period oscillations of bacteria, while shorter oscillations suppressed SMDD neurons (**Fig. 2e,f**). These data are consistent with previous studies showing a role for SMD neurons in biasing head movements and locomotion [19,39], likely altering the trajectory of movement in gradients of bacterial stimuli in three dimensions. Collectively, these studies identify the patterns of bacterial stimuli that drive maximal responses in the sensory ON and OFF, AVA, RME, SMDD and SMDV clusters.

### *C. elegans* neural activity can be embedded in hyperbolic space

To further analyze the coordination between different neural activity patterns, we attempted to reduce the dimensionality of the data. Previous studies have used Principal Component Analysis and showed that changes in neural activity can be analyzed by evaluating the first three principal components [19]. However, we found that the first three components could only account for about 37% of the variance in our data (**Fig. 3a, S3a**). Given that activity of the *C. elegans* head neurons can be hierarchically organized [41,42,43], and that hierarchical organization is generally expected based on efficiency principles [44], we sought to determine whether a better dimensionality reduction can be obtained with a hyperbolic metric that approximates hierarchical processes [45]. We used a Bayesian Hyperbolic Multidimensional Scaling to embed our neural activity data in hyperbolic space [46]. Hyperbolic embedding made it possible to better maintain the original distances between the cells (**Fig. 3d** vs. **Fig. 3b**).

**Figure 3.**
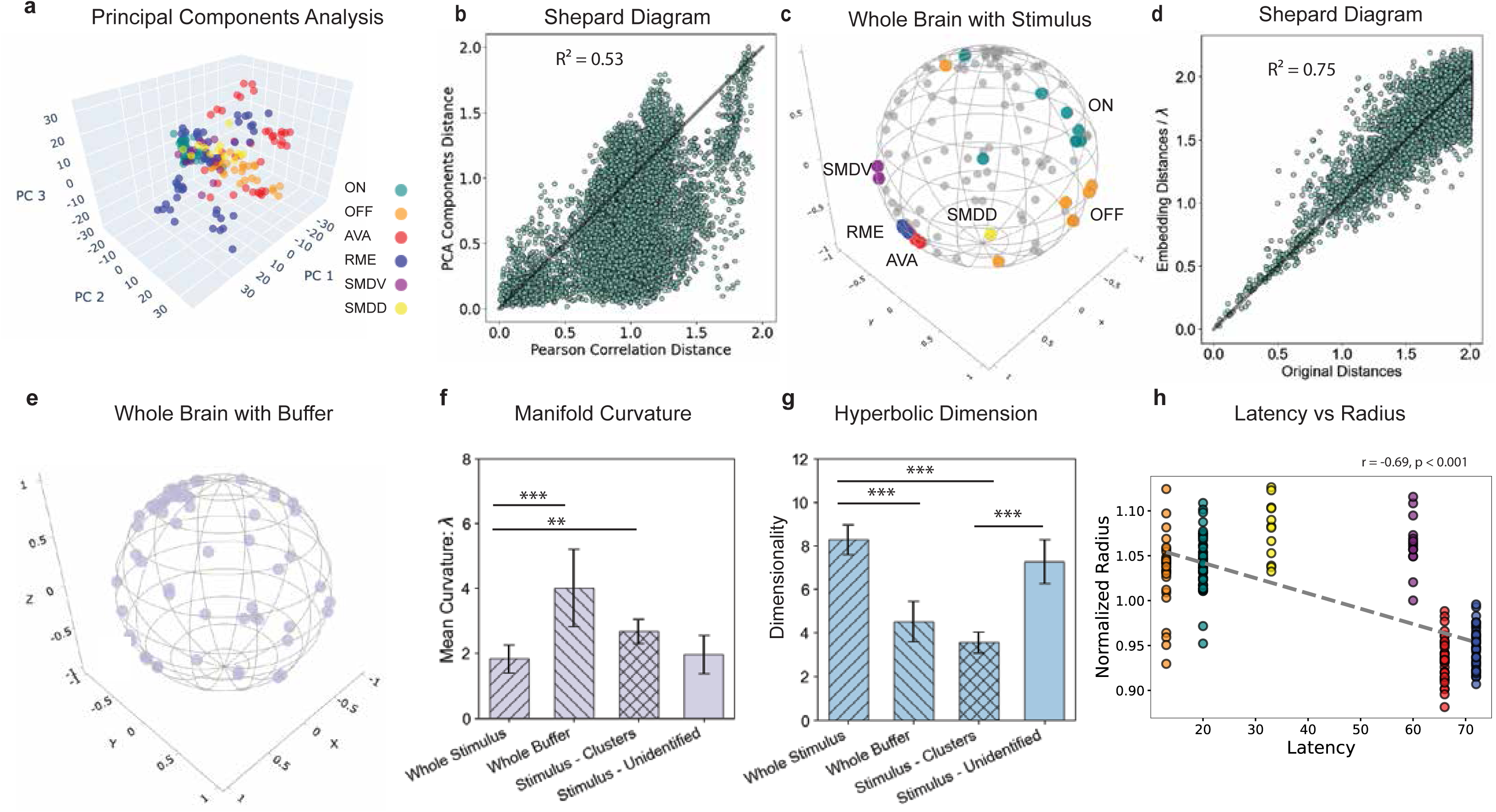
*C. elegans* neural activity can be embedded in hyperbolic space. (**a**) Principal Components Analysis (PCA) showing the top three principal components of all clustered cells from seven animals exposed to stimulus. (**b**) Shepard diagram depicting the relationship between original pairwise distances between calcium traces of neurons. Original distances were defined as two times one minus the squared Pearson correlation, 𝑑_𝑖𝑗_ = 2(1 − 𝑟^2^), PCA distances were calculated based on the top three principal components. (**c**) Poincaré disk representation of a hyperbolic embedding for a single animal exposed to stimulus. Clustered cells are denoted by color and labeled, while unclustered cells are shown in grey. (**d**) Shepard diagram comparing original pairwise distances and distances reconstructed from the hyperbolic embedding normalized by curvature. Original distances were defined as 𝑑_𝑖𝑗_ = 2(1 − 𝑟^2^). (**e**) Poincaré disk representation of a hyperbolic embedding of the whole brain from a single animal exposed to buffer only. (**f**) Comparison of mean curvature (λ) values across different groups. Bars represent the mean ± standard deviation (SD) for each group. Groups include “Whole Stimulus,” “Whole Buffer,” “Stimulus – Clusters,” and “Stimulus – Unidentified.” (**g**) Dimensionality of different datasets determined by Bayesian Information Criterion (BIC) analysis, shown as sample mean ± SD. (**h**) Scatter plot of normalized hyperbolic radius versus response latency across all worms exposed to stimulus. Each point represents a clustered cell, colored by cell type (from left to right: ON sensory, OFF sensory, SMDD, SMDV, AVA, RME). The dashed black line shows the best-fit linear regression using ordinary least squares. The Pearson correlation coefficient, r, quantifies the strength and direction of the linear association. For panels **f** and **g**, statistical significance was assessed using independent two-sample *t*-tests assuming unequal variance (Welch’s *t*-test). For panels **b** and **d**, R^2^ indicates the goodness-of-fit between original and embedded distances, with R^2^ =1 denoting perfect preservation of distances. In all panels, only significant differences are annotated. \**P <* 0.05; \*\**P <* 0.01; \*\*\**P <* 0.001; \*\*\*\**P <* 0.0001.

To visualize neural activity in a hyperbolic embedding, we used the Poincare ball visualization (**Fig. 3c**). This is a compact representation of a n-dimensional hyperbolic space, in which all the space is mapped into the interior of a unit sphere [45]. We found that neurons belonging to the AVA and RME clusters were close to one another, while those in other clusters were separated. Next, we found that the neural activity of a single animal experiencing a stimulus-to-buffer sequence was 8 dimensional, based on Bayesian Information Criteria (**Fig. 3g**, see methods). Neural activity from animals experiencing buffer-to-buffer sequences without any bacterial stimuli (**Fig. 3e**) had a lower dimensionality, with four dimensions, on average (**Fig. 3g**). This decrease in dimensionality was mostly due to the unclustered cells that were active only in the stimulus-to-buffer data but not in the buffer-to-buffer case (**Fig. 3g**).

To further test how strongly neural embedding differed from the Euclidean space, we measured the curvature, -λ^2^, of the space occupied by the data (**Fig. 3f**, see methods). We found that scale of the curvature, λ was significantly greater than 0 in all the four datasets (stimulus-buffer, buffer-buffer, clustered cells under stimulus and unidentifiable cells under stimulus). Because λ was at least 2, the space can be considered strongly hyperbolic. We note that when comparing curvature between stimulus-buffer and buffer-to-buffer conditions, one should consider that buffer-to-buffer conditions was lower dimensional. Therefore, curvature cannot be directly compared between these two cases.

There was a systematic relationship between neuronal type and position within the hyperbolic space. The interneuron clusters AVA and RME had more complex responses and took more central positions in the embedding (**Fig. 3h**). This positioning indicates their hub-like properties within the network [23]. By comparison, motor neurons (SMDD and SMDV) and sensory ON/OFF neurons took more peripheral positions (**Fig. 3h**). We also found that there was a negative relationship between response latency and a cell’s radius in the hyperbolic embedding. The AVA and RME clusters took the longest to respond to bacteria and had smaller radii when compared to ON and OFF sensory clusters. Collectively, this data suggest that the animal movement is controlled by the AVA and RME clusters. These neurons drive activity in the SMDD and SMDV neurons. At the same time, activity of AVA/RME interneurons can be biased based on sensory input from ON/OFF neurons to allow for adaptation to distinct states.

### Three dimensions are sufficient to distinguish all six neuronal clusters

Although whole brain activity required eight hyperbolic dimensions, the activity of clustered neurons could be described with just three hyperbolic dimensions (**Fig. 3g**). This makes it possible to visualize their activity (**Fig. 4a,b**, see methods). One of the three dimensions separate sensory ON/OFF neurons (**Fig 4a, b**). Another dimension separated the SMDD and SMDV clusters (**Fig. 4c,d**), a dorsal/ventral dimension. Initially, we did not observe a clear separation of the AVA and RME clusters (**Fig. 4e,f**). However, in our experiments, each animal experienced one of two different patterns of bacterial stimuli (see methods). When the data was separated according to the stimulus pattern, we did observe a separation of the cells between the AVA and RME clusters (**Fig. 4g**). We also mapped the AVA and RME cells in tangent space and observed that a single dimension could separate their activity (**Fig. S4a**). Finally, we also combined all the AVA and RME cells from separate embeddings for each animal onto a single Poincare ball and confirmed separation of the two clusters (**Fig. 4h**), which corresponds to a forward/backward motion dimension. Collectively, these data suggest that neurons in the six identified clusters of neurons in animals experiencing bacteria-to-buffer patterns can be discriminated using the ON/OFF, forward/backward, and dorsal/ventral dimension.

**Figure 4.**
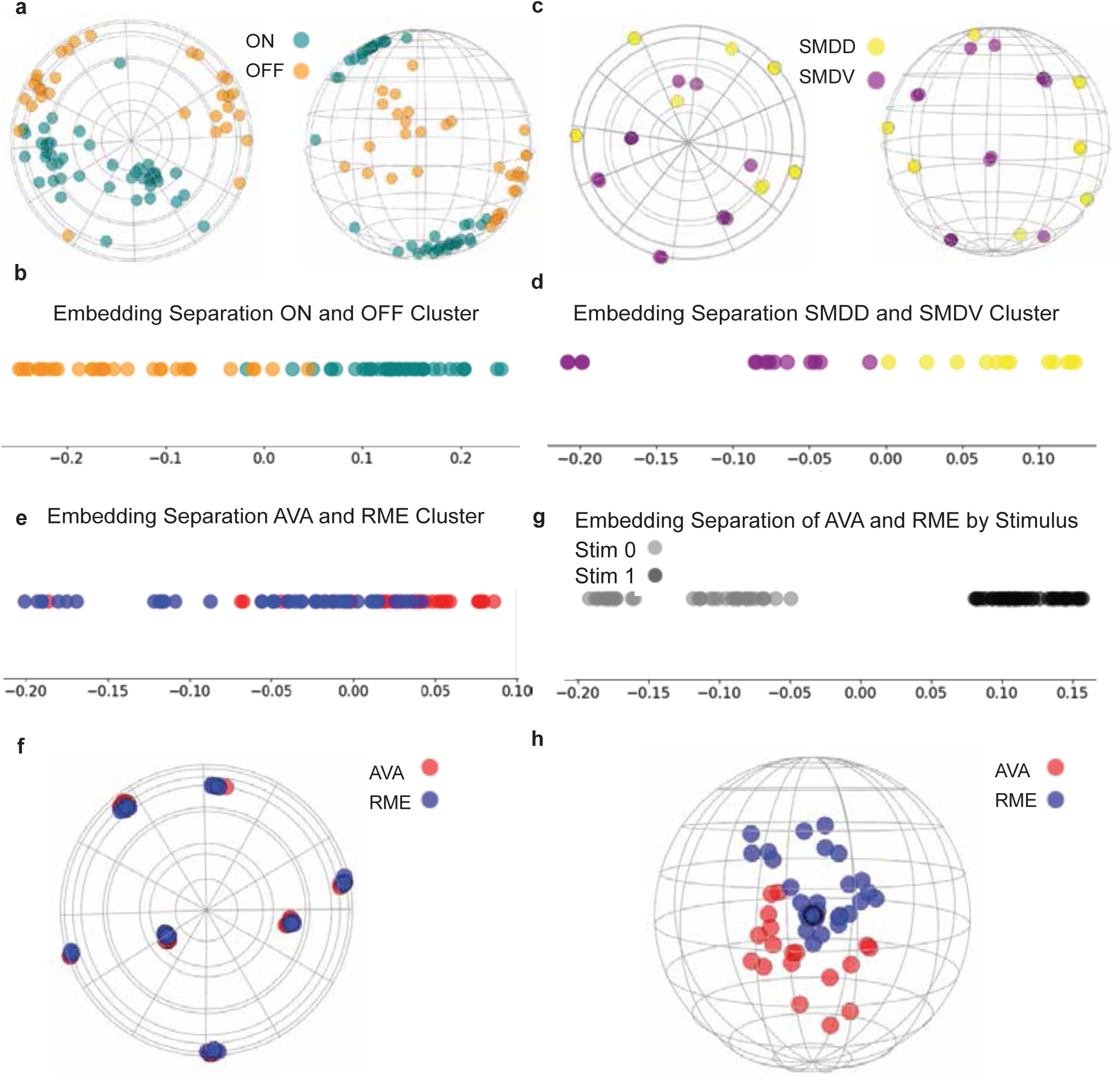
Three dimensions are sufficient to distinguish all six neuronal clusters. (**a**) Two views of Poincaré disk, visualizing a 3-dimensional hyperbolic embedding of all clustered ON/OFF sensory cells in animals exposed to patterns of bacterial stimuli. (**b**) 1-dimensional scatter plot of ON/OFF sensory cells depicted in Poincaré disk, colored by cluster identity. (**c**) Two views of Poincaré disk, visualizing a 3-dimensional hyperbolic embedding of all clustered SMDD and SMDV cells in stimulus condition worms, colored by cluster identity. (**d**) 1-dimensional scatter plot of SMDD and SMDV cells depicted in Poincaré disk. (**e**) 1-dimensional scatter plot of AVA and RME cells across all 7 stimulus condition worms. (**f**) Poincaré disk representation of all AVA and RME cells across all 7 stimulus condition worms. (**g**) 1-dimensional scatter plot of AVA and RME cells across all 7 stimulus condition worms, colored by stimulus pattern. (**h**) AVA and RME cells from each worm, individually embedded into hyperbolic space, and plotted altogether in a Poincaré disk.

## Discussion

In this work we show that *C. elegans* head neurons can be classified into at least six clusters (ON and OFF sensory, AVA, RME, SMDV, and SMDD) based on their responses to patterns of bacterial stimuli. We also defined the bacterial stimulus pattern that drives maximal responses in each of these six clusters. We also obtained evidence that this neural network is likely to be hierarchical by using a hyperbolic geometry to embed the neural activity data. Notably, we find that the dimensionality of the data in animals experiencing bacterial stimuli was twice compared to those exposed to control buffer stimuli. Moreover, this additional dimensionality observed in the stimulus condition is driven by neurons whose activity does not correlate with any of the six identified clusters. Finally, we show that this 8-dimensional network can be further reduced to three (sensory, forward-backward, and dorsal-ventral) along which the six neuronal clusters can be separated. Our study provides a framework for analyzing high-dimensional neuronal activity patterns linking sensory responses to control interneurons to motor neurons.

Our strategy of monitoring changes in neural activity patterns in animals experiencing patterns of bacterial stimuli has enabled us to define the relevant pattern of stimuli that drives maximal response in each identified cluster. This pattern of bacterial stimuli is analogous to studies identifying the receptive field of a neuron. In the visual system, these receptive fields have revealed how visual information is integrated across different cell types and how complex features of the visual stimulus emerge by combining local features [47,48,49]. We show that the maximal responses of ON and OFF sensory neurons occur within 15 seconds of a rapid increase and decrease in bacterial stimuli respectively. This time window is consistent with previous studies showing that sensory neurons have their maximal responses within 15 seconds of addition or removal of bacterial stimuli around the animal’s nose [35,36]. Although we and these previous studies find that activity in sensory neurons changes within 1 second of change in bacterial stimulus around the animal nose, the 15-second time window for maximal response might represent cellular calcium and GCaMP dynamics [50] within the sensory neurons. In contrast, we find that changes in bacterial stimuli many tens of seconds prior to the maximal responses in maximal responses in AVA, RME, SMDD and SMDV neurons. Prior studies have found that AVA neurons are active during reverse, while RME neurons are active during forward movement [19,25]. Additionally, SMDD and SMDV neurons bias the movement of the head towards dorsal and ventral directions respectively [30,31]. This relatively long-time window for change in bacterial stimuli is unexpected and might represent dynamics in cellular calcium, GCaMP signals along with information processing within the densely interconnected *C. elegans* network [18,31]. The feature selectivity of AVA and RME neurons was more complex than that of either the sensory ON/OFF neurons or the motor SMDD/SMDV neurons. Specifically, it required at least two stimulus features for their description, one excitatory and one suppressive (**Fig S2g,h**). This type of selectivity is well documented for neurons in other species and suggests the classical complex cells from the primary visual cortex [51]. These cells often have excitatory and suppressive components [52, 53].

Our approach of using hyperbolic embedding is designed to capture hierarchical aspects of neural activity [32,33,34,35]. Previous studies have shown that the *C. elegans* neural network is hierarchical when analyzed using neural activity patterns or cellular anatomy underlying behavior [41]. In a hierarchical network, the distance between two points, A and B, is evaluated as the distance between point A, to the first common ancestor of these two points and then the distance to point B. The shortest distance between two points in a curved space, otherwise known as the geodesic [54], is thus represented differently than the shortest distance between points in a flat space, using Euclidean distance measures. Consequently, distances between neurons in a curved space in a hierarchical network are represented differently than in a flat space [22]. Here we demonstrate that the whole brain when exposed to chemosensory stimuli shows an increase in its hyperbolic dimensionality. We interpret the increase in dimensionality of the corresponding neural network in animals experiencing bacterial stimuli compared to their control buffer as an increase in the hierarchical nature of the cell network as it is represented by the calcium time series of each individual neuron. Moreover, we also find that the latency between change in bacterial stimuli and maximal response of a neuronal cluster also increased with its hyperbolic radius, implying that the cluster’s preferred stimulus feature and its relationship to the bacterial stimuli was a leading contributor to the hierarchy of the neural network. Hierarchical analysis such as this has been found to aid in interpretation of neuronal activity clusters [32, 33]. Furthermore, we find that in hyperbolic space, three dimensions can fully separate the six clustered cells. These include the sensory stimulus (ON or OFF), the backward and forward (AVA and RME), and dorsal-ventral (SMDD and SMDV) dimensions. We suggest that additional features including the responses of unclustered neurons and their relationship with bacterial stimuli might contribute to the additional dimensionality of eight. Collectively, our approach defines six clusters of *C. elegans* neurons and identifies hyperbolic geometry as well suited to represent neural activity data. More broadly, we reveal how responses in each neuronal cluster are related to sensory stimuli and how the relationship between these clusters is altered by sensory input, key steps in relating sensory input to behavioral outputs.

<u>Speculation:</u> We suggest that bacterial stimulus drives responses within *C. elegans* neurons, which are organized in a hierarchical fashion, like what has been previously observed [41]. While a similar hierarchy is also observed when animals experience changes in buffer solution around their nose, their dimensionality is much reduced. Moreover, we suggest that the AVA and RME neurons serve as network hubs that affect SMDD and SMDV nodes. These four clusters might represent the basal state of the animal, which is modified by the sensory input. We further speculate the long time-window between changes in bacterial stimuli and maximal responses in AVA, RME, SMDD, and SMDV clusters is correlated with time required to process sensory information in this hierarchical network.

## Methods

### Head neuronal imaging

We used two transgenic strains that expressed GCaMP. The primary strain (ZIM294) expressed GCaMP5K in the nuclei of all neurons (*mzmEx199* [*Punc-31::NLSGCaMP5K*; *Punc-122::GFP*]). To identify neurons associated with activity patterns observed in ZIM294, we used a strain (OH15500) that expressed GCaMP6s and NeuroPAL (*otIs669*[*NeuroPAL*];*otIs672*[*Panneuronal GCaMP6s*]). Cells were identified according to the map described by Yemini and colleagues [27]. We monitored changes in GCaMP fluorescence using a Zeiss LSM 880 with Airyscan. Acquisition was done in 2-micron z-steps. In ‘Fast’ mode, the Airyscan images the entire head of the adult worm at about 1.5 volumes per second. Worms were typically imaged for approximately ten minutes. We then used piecewise rigid registration to remove motion artefacts [55] and non-negative matrix factorization to isolate individual neurons and extract their fluorescence values [56]. Out of a total 189 neurons in the head, our approach identified 50-100 neurons per animal.

### Stimulus delivery

Day 1 adult animals were washed in M9 and loaded into a microfluidic device that trapped the worm body while exposing only the nose to stimulus flows [25]. Animals were also treated with 1.5 mM of the paralytic tetramisole hydrochloride to suppress most perceivable worm movement. The movement of untreated worms proved too difficult to motion correct. We delivered precise patterns of fluctuating bacteria and M9 buffer liquid flows using an Arduino to send pulses to a valve controller. The bacteria solution was prepared as a 1:1 resuspension of a bacterial culture (OD600 = 0.4) in M9 buffer as previously described [36]. The controller determines whether bacteria or buffer is routed to the nose of the trapped worm. Worms were exposed to binary patterns of bacteria and buffer. Several different stimulus protocols are used in this study. In the base protocol, the trial is divided up into pulse blocks of ∼15 seconds. The pattern is constructed using transition probabilities: *p*(switch on | off) = 0.2 and *p*(switch off | on) = 0.4. In the faster protocols, the same switch probabilities are used but the pulse blocks have length ∼1.5 seconds. Buffer data in this study was obtained from animals experiencing switches in the M9 buffer flow around their nose [26].

### Classification of neurons

We identified primary sensory neurons by looking for neurons that were either positively correlated (ON cells) or negatively correlated (OFF cells) with bacteria pulse onset during bacteria-to-buffer stimulus sequences. ON cells were categorized as those neurons that immediately increased activity upon all bacteria onsets and immediately decreased upon bacteria offsets. OFF cells were classified as neurons whose activity decreased upon bacteria onsets and increased upon bacteria offsets. Within each animal, the cells whose traces had >85% Pearson correlation to the classified ON and OFF representative cells were also labeled as ON and OFF cells, respectively.

We also observed a pair of anti-correlated and bistable neurons and a second pair of anti-correlated and moderately fast neurons. We used the NeuroPAL–GCaMP6s strain to identify these neurons and determined the anti-correlated and bistable neurons to be RME motor neurons and AVA command neurons, and the second anti-correlated pair with faster dynamics to be SMDD and SMDV. Using AVA, RME, SMDD, and SMDV as representative neurons, we sorted non-sensory neurons into clusters based on how their activity correlated with these four representative neuron pairs. AVA and RME were previously shown to have strong positional stereotypy and thus, after identifying these neurons with NeuroPAL, their activity signature and stereotyped location made them easy to identify in the absence of NeuroPAL. We classified the remainder of the cells whose traces had a strong Pearson correlation (>85%) with the identified RME neurons or AVA neurons. SMDV and SMDD were often the sole members of clusters within each organism. We were able to subsequently identify SMDV and SMDD neuron clusters in non-NeuroPAL-GCaMP5K animals and classified the remainder of the cells whose traces had a strong Pearson correlation (>85%).

### Maximum noise entropy analysis

The probability of observing a response in a neuron pertaining to a specific neural cluster was defined by a nonlinear function of the stimulus vector, 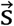.

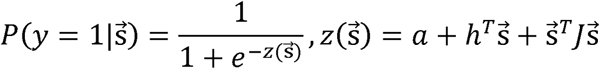

Upon training the model, we extracted the low rank matrix 𝐽 which was determined by minimizing the negative log-likelihood [33,34]. We proceeded to diagonalize 𝐽 which gave us the receptive field. As input, we used a stimulus pattern, 𝑠,for each corresponding response, 𝑦. For each cell in our analysis, the calcium response for a given time bin was averaged and paired with a stimulus vector of averaged time bins of stimulus that preceded this response. For each cell 1000 time points were analyzed. Training was done by splitting the calcium time series data into 4 separate jackknives. There were four training iterations, each with 25% of the data (one jackknife) omitted each time, and used as the cross validation (CV) set, and the other 75% (three jackknives) used as the training set. The jackknife used for CV was alternated each iteration. The 𝐽 matrix was obtained four times, once for each iteration of training. These four 𝐽 matrices were averaged, the resulting matrix was decomposed, and its top eigenvalue’s corresponding eigenvector was taken and flattened into a matrix. The resulting flattened matrix was then factorized and the top eigenvalue by magnitude was selected, its eigenvector was the receptive field, RF.

### Maximum noise entropy data processing

For training and subsequent model weight extraction, data was binned into stimulus-response pairs. Every 𝑦 and 𝑠 pair, served as the label and data pairs during model training. The dimensionality of the input layer was determined by the dimensionality of the input stimulus vector 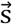. For the entire training set: y = y an average across several subsequent points 𝑛 in the normalized calcium response time series, starting at t_0_.

Stimulus was a vector, 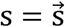, defined as 𝑛 number of bins of 𝑥 data points in each bin the stimulus time series, starting at t_0_ and going back in time.

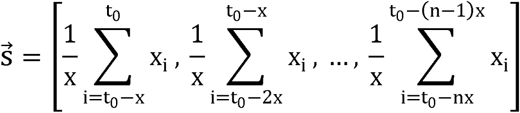

For each neuron type, the model was run 4 times. The data and corresponding labels were divided into 4 jackknives. Three jack knives were used for training, and one was left out for cross-validation. A total of four sets of model weights were collected. Given four symmetric matrices, J_0_, J_1_, J_2_, J_3_ each was flattened into a vector:

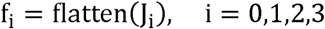

We computed the combined outer product matrix 𝐴 by summing the outer products of the flattened vectors:

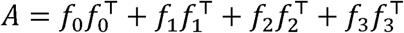

And then we symmetrized matrix 𝐴_𝑠_:

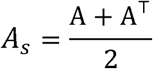

We performed Eigen Decomposition on the symmetrized matrix. Then we selected the eigenvector corresponding to the largest eigenvalue where:

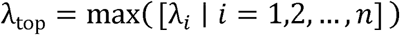

We reshaped the Top Eigenvector into a matrix, symmetrized the reshaped matrix, and performed Eigen Decomposition on the symmetrized matrix. Finally, we select the eigenvector corresponding to the largest eigenvalue. RFs were determined by selecting the corresponding eigenvector for the top in magnitude eigenvalue.

### Stimulus Component Nonlinearity

We computed the dot product between the RF vector found earlier using MNE and the stimulus vectors. This projection of the stimuli onto the RF provided us with a component of the stimulus. All stimulus vectors for each neuronal cluster were projected onto that cluster’s RF. As we plotted the response of the neurons as a function of the resulting dot products, we observed a nonlinearity for each neuronal type. The same operation was done with all stimulus vectors and the eigenvector that corresponded to the second largest in magnitude eigenvalue of the processed 𝐽 matrix obtained from MNE. This provided the top two components of the stimulus. Calcium response for each type of neuron as a function of Component 1 and Component 2 was plotted as heatmaps. Individual components were plotted as 2D nonlinear gain functions of the stimulus along the diagonal margins of each heatmap. These represented the response as a function of the projection of the stimulus onto the filter. Each component was normalized to have zero mean and unit variance.

### Bayesian Hyperbolic multidimensional scaling

Bayesian Hyperbolic Multidimensional Scaling (BHMDS) [46] was used to calculate the curvature of the space, λ, and the Bayesian Information Criteria (BIC) in normalized calcium trace whole brain data. The given dissimilarities matrix δ_ij_ can otherwise be modeled by dividing a distance matrix d_ij_ by a scale of curvature parameter, λ, and adding Gaussian noise.

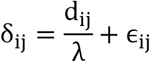

Given the dissimilarities matrix, B-HMDS seeks to find the set of embedding points 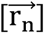 that minimizes the stress function, thereby minimizing the distance between the given dissimilarities and the distance matrix d_ij_.

In instances where cell counts from animal to animal were not equivalent, up-sampling was performed on the low cell count animals, to match the cell numbers of the highest animal. Cells from the low population condition animals were randomly chosen, and included twice, to ensure they were not exact copies of pre-existing cell traces, thus their distances would not be 0 (the same cell) in the dissimilarities distance matrix.

Points in hyperbolic native space, 𝑟, were computed according to the law of cosines, and 𝑙, the distance between points can be computed like so, where λ helps define the curvature of the space.

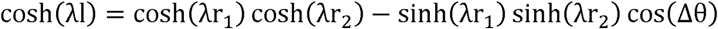

Curvature of the space, can be modeled as:

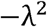

Where λ, is the factor that determines the scale of the curvature of the space.

### Bayesian Information Criteria

The probability of observing the given dissimilarities matrix conditioned of a given dimensionality of hyperbolic space is given by:

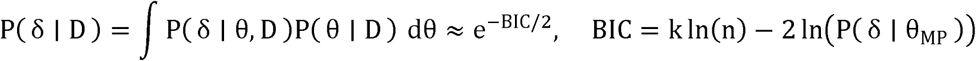

This formula calculates the highest likelihood of seeing the given dissimilarities distance matrix given the smallest number of k parameters, the smallest dimensionality.

### Dimensionality analysis of data in hyperbolic space

We plotted neurons from all seven stimulus-buffer organisms in a Poincare ball. Their embedding coordinates were plotted in the Poincare representation of hyperbolic space. We used a logarithmic map to map each space in the embedding to its corresponding point in the Euclidean tangent space at the origin. We defined the average position of each category in the tangent space, and created a unit vector in the tangent space by calculating the normalized difference in means between the two-point clusters. We proceeded to project each individual cell’s coordinates onto this dimension.

## Acknowledgements

We thank Manuel Zimmer, Oliver Hobert and CGC (Caenorhabditis Genetics Center) for worm strains and Uri Manor, Tong Zhang and the Waitt Advanced Biophotonics Center for advice and assistance with our imaging experiments. We also thank Ryan Rowekamp, Anoop Praturu, Haodong Qin, Javier How, for discussions and comments on the manuscript. This work was supported by grants from the National Institutes of Health NIH 5R01MH096881 (S.H.C); Esther A. & Joseph Klingenstein Fund, Simons Foundation, and Hypothesis Fund(E.Y); U19NS112959, P30AG068635, AHA-Allen Initiative in Brain Health and Cognitive Impairment award 19PABH134610000, National Science Foundation (NSF) grants IIS-1724421, PHY-4213080, NSF Next Generation Networks for Neuroscience Program award 2014217 and by the Edwin Hunter Chair in Computational Neuroscience (T.O.S.). The funders had no role in study design, data collection and analysis, decision to publish, or preparation of the manuscript. I.R. was also supported by the Pathways in Biological Sciences (PiBS) Fellowship. Z.T.C. was supported by a fellowship from the Dan and Marina Lewis Foundation. K.T.Q. was supported by a Postdoctoral fellowship from the Paul F. Glenn Foundation.

## Author Contributions

I.R. developed computational methods and analyzed data, Z.T.C. designed experiments. Z.T.C. and J.J.H. performed experiments. E.Y. contributed provided strains. I.R., Z.T.C., and K.T.Q. analyzed data. I.R., T.O.S. and S.H.C. wrote the paper. T.O.S. and S.H.C. supervised research and obtained funding for the project.

---

**Figure S1.**
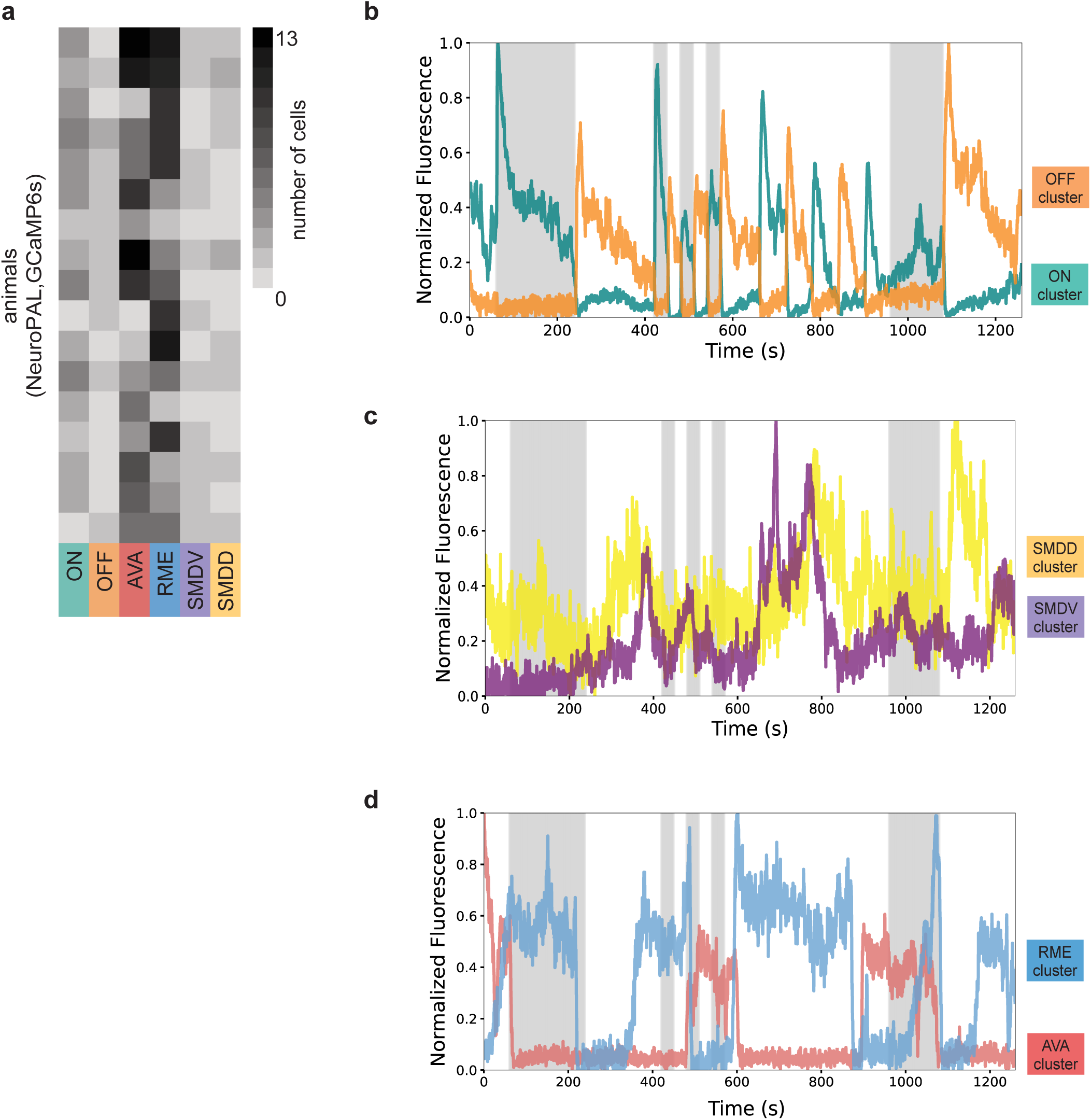
Calcium responses in animals exposed to buffer alone. (**a**) NeuroPAL-GCaMP6s calcium activity. (**b**) Representative ON/OFF sensory cells in buffer-to-buffer session. (**c**) Representative neurons in buffer-to-buffer session with highest Pearson-correlation to SMDD and SMDV neurons found in stimulus-to-buffer session. (**d**) Representative neurons in buffer-to-buffer session with highest Pearson-correlation to RME and AVA neurons found in stimulus-to-buffer session.

**Figure S2.**
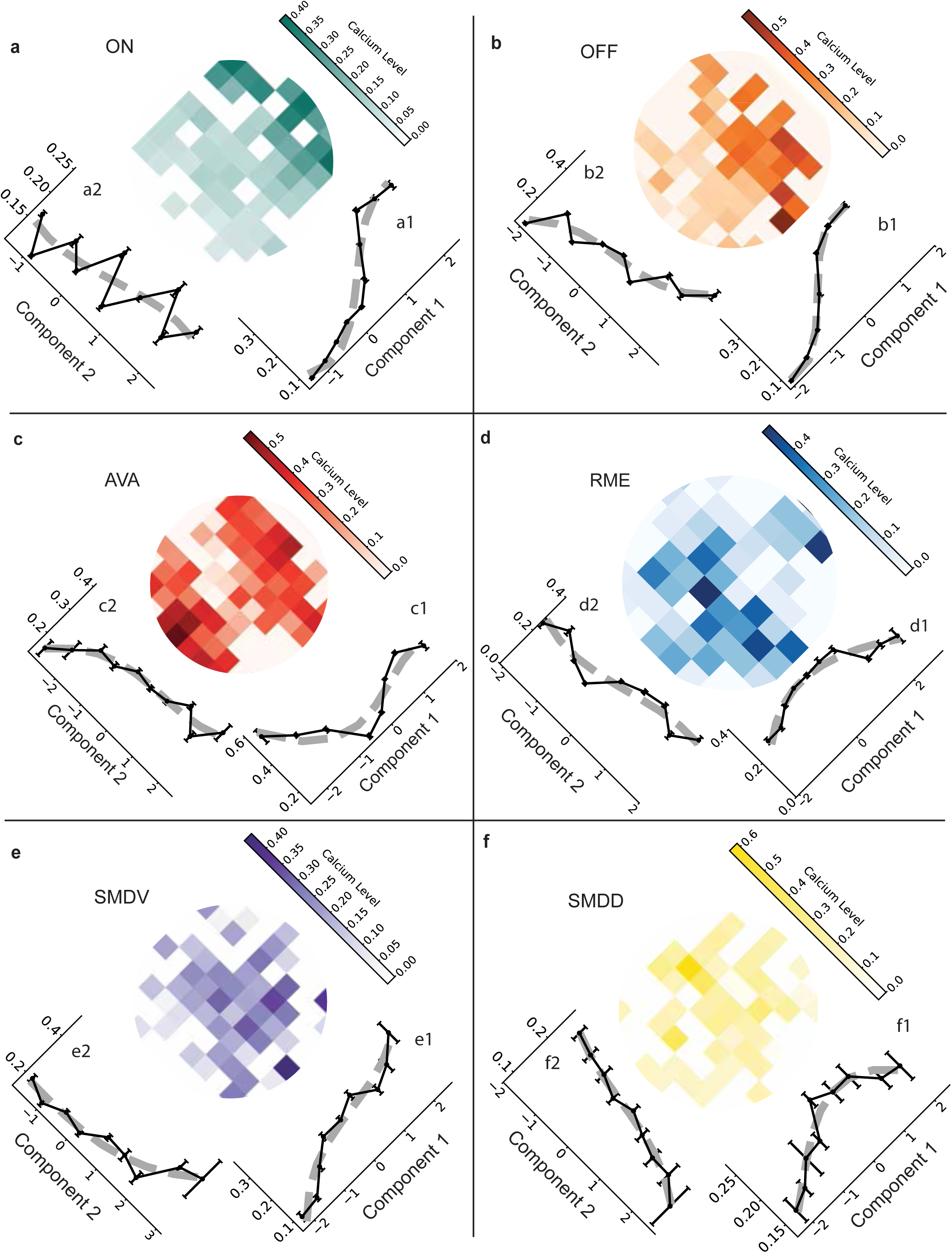
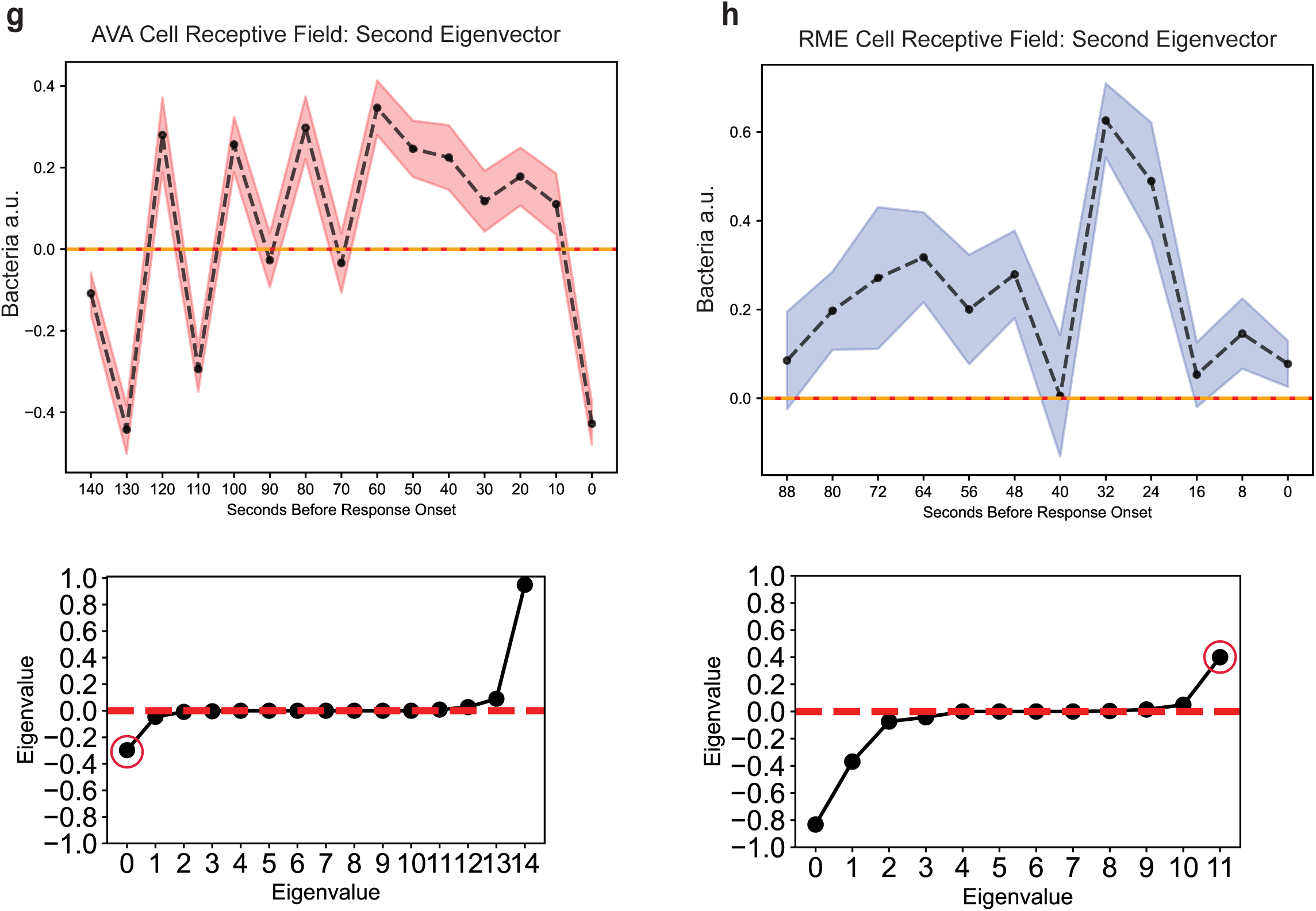
Stimulus components altering calcium responses across the six neuronal clusters. (**a**-**f**) Effects of stimulus components on calcium responses across six clusters of neurons. The first and second stimulus components are shown individually in linear side plots. Stimulus components, indicated by black lines, were normalized to unit variance and plotted in units of standard deviation. Grey dashed lines represent polynomial fits. Central heatmaps in each subpanel display normalized calcium levels as a function of the two stimulus components. (G–H) Secondary receptive fields for AVA and RME neurons. The second-highest eigenvector is plotted as a black dashed line, corresponding to the second-largest eigenvalue (in absolute value). The shaded colored region in each plot represents the standard error of the mean (SEM) across 20 bootstrap samples. Eigenvalue decompositions for each respective cell time series are shown below the corresponding eigenvector plots, with the second eigenvalue circled.

**Figure S3.**
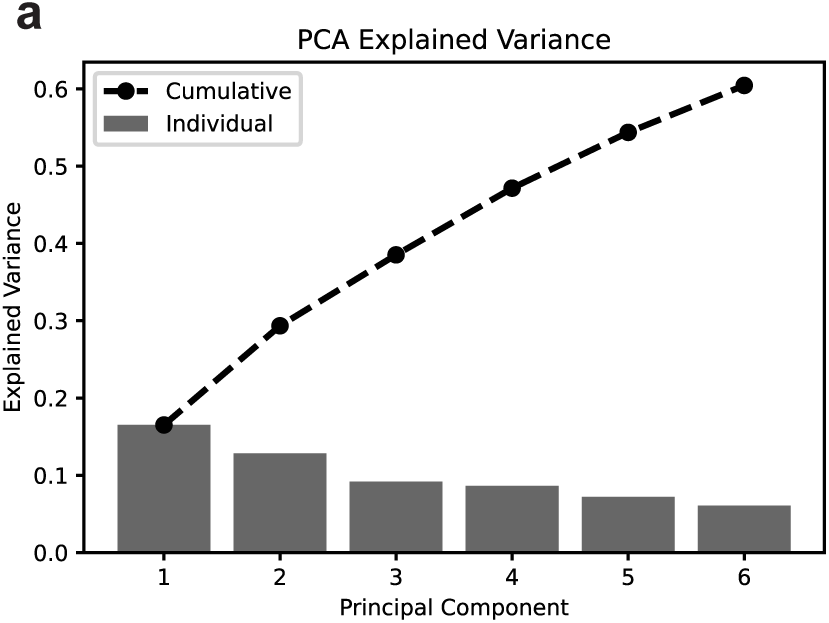
Principal components affecting variance in calcium data. (**a**) Principal component analysis (PCA) variance explained by each of the first six principal components. Bars represent the proportion of variance explained by each individual component, while the dashed line shows the cumulative variance explained.

**Figure S4.**
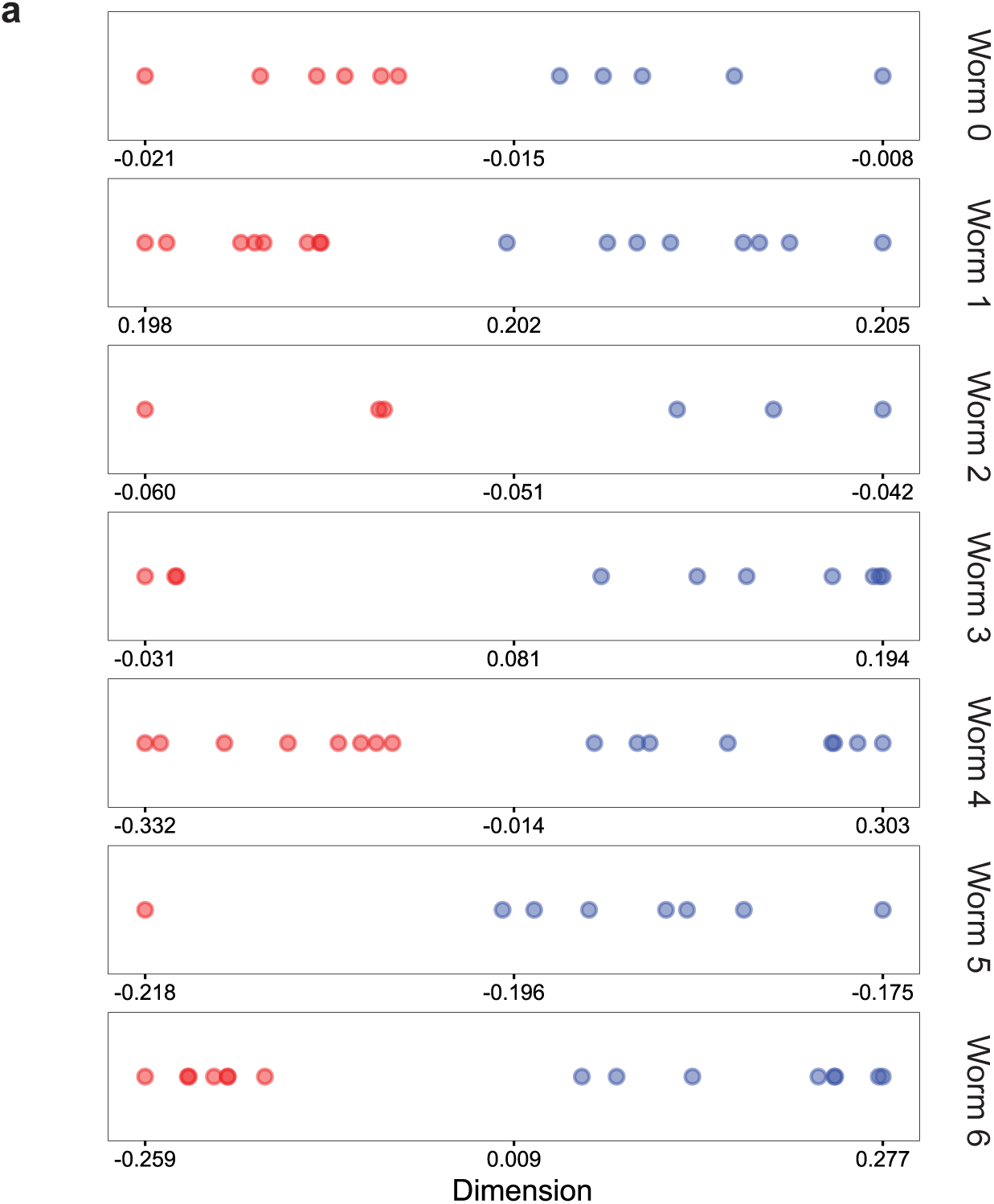
Properties of AVA and RME cells embedded in hyperbolic space. (**a**) 1-dimensional plots of AVA and RME cells embedded individually into hyperbolic space, colored by cell type.

