## Supplementary figures and images for "A framework for analyzing *C. elegans* neural activity using multi-dimensional hyperbolic embedding"

### Supplementary FIgure 1

##### Supplementary 1S

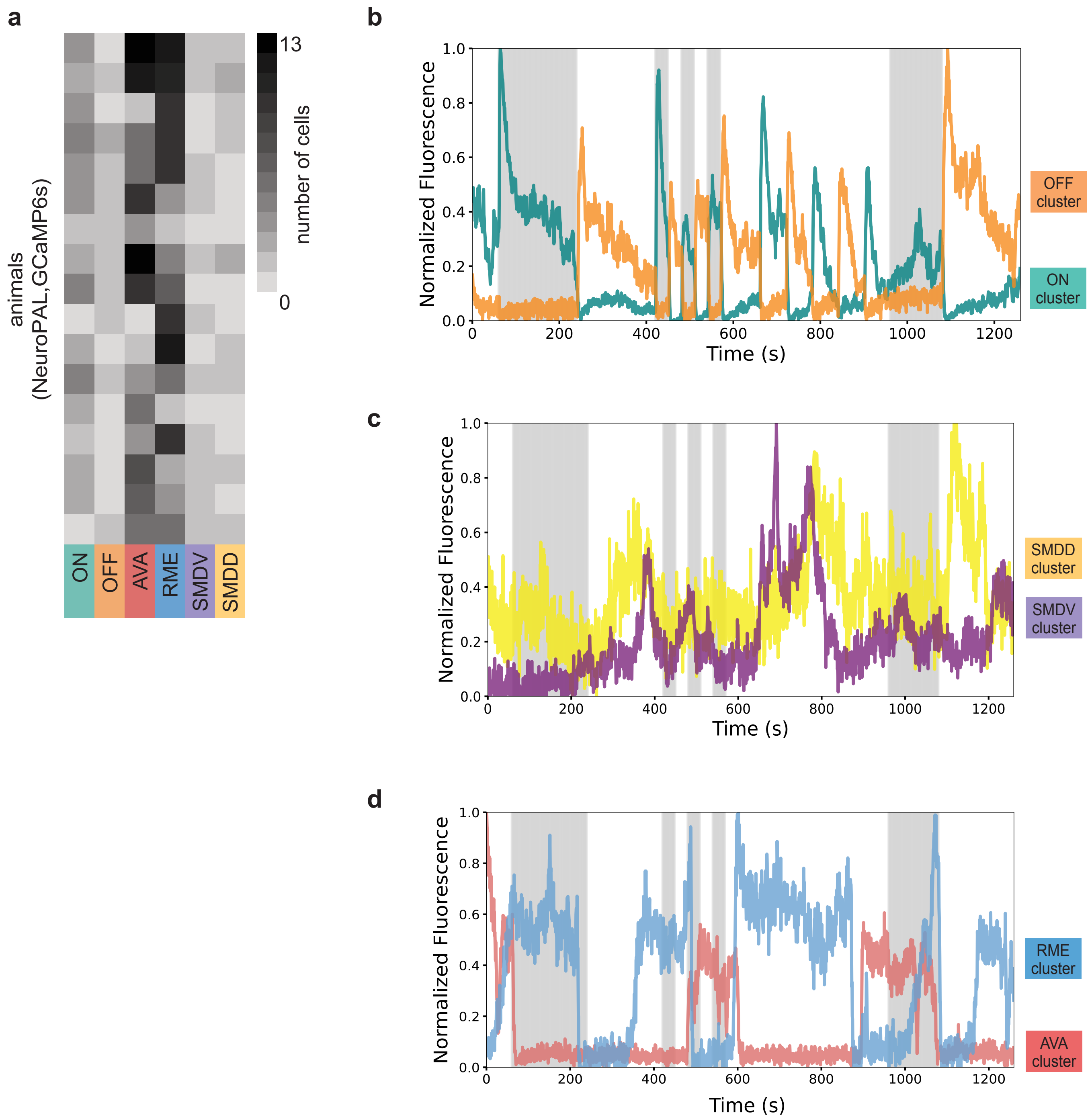

### Supplementary Figure 2

Supplementary 2S

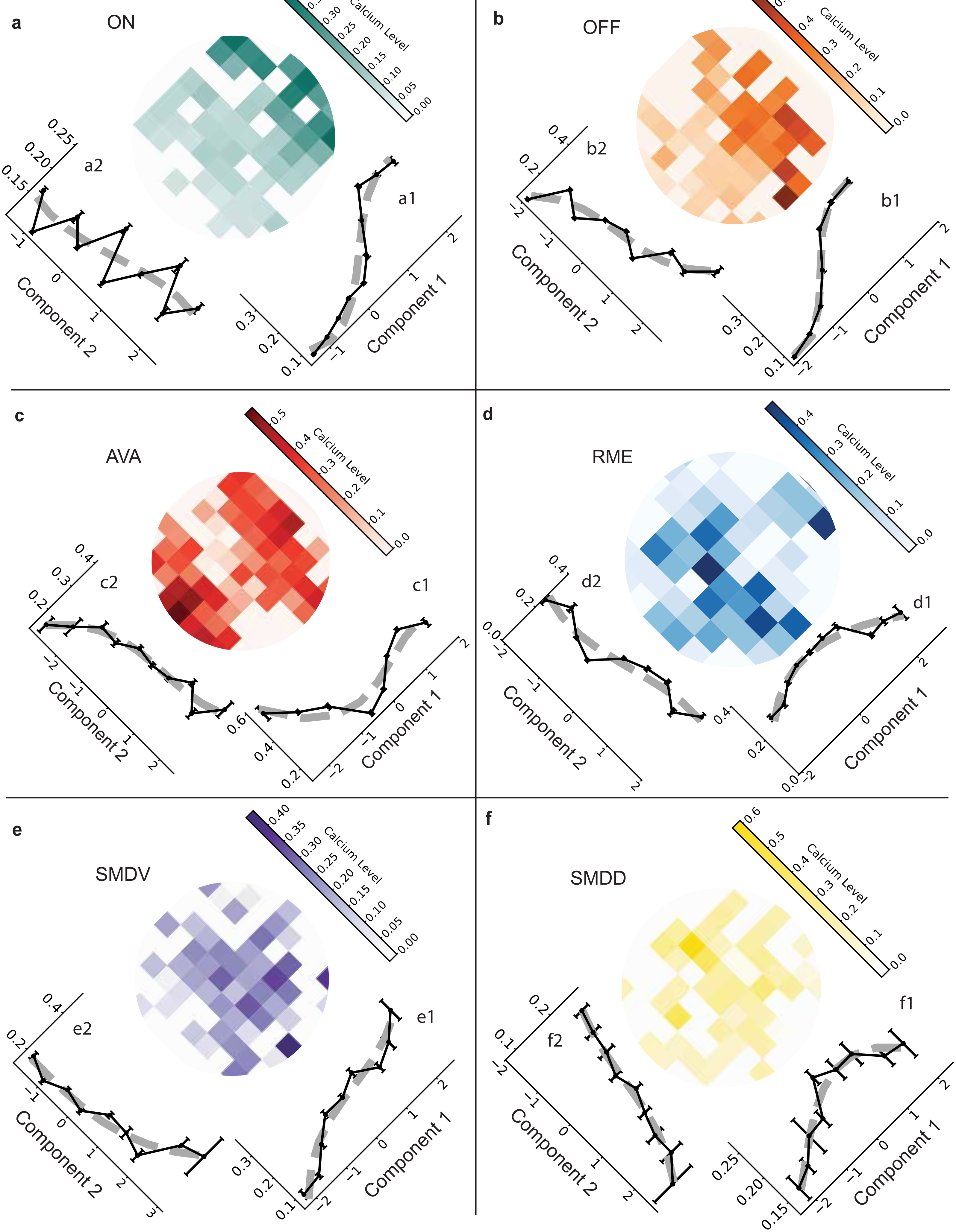

### Supplementary Figure 3

#### Supplementary 3S

**a**

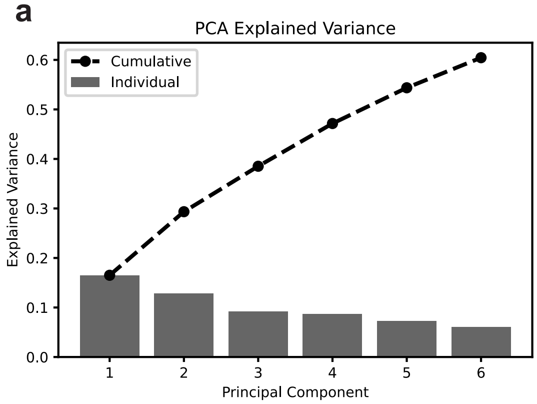

### Supplementary Figure 4

##### Supplementary 4S

**a**

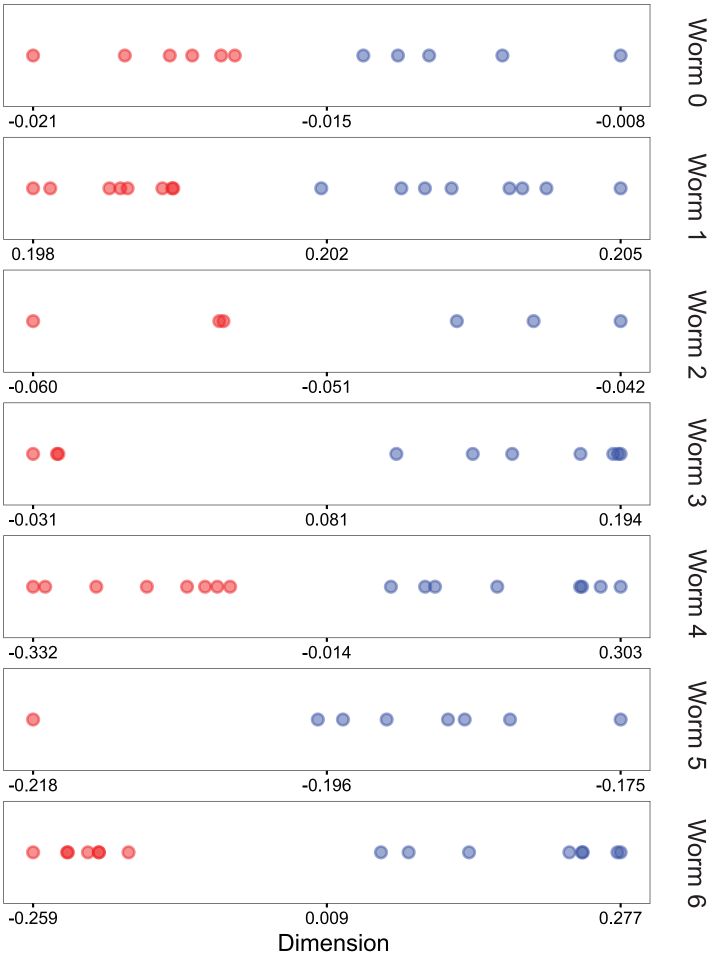
